# Optogenetic activation of liver-innervating vagal sensory neurons increases anxiety-like behavior in mice

**DOI:** 10.1101/2025.08.04.668542

**Authors:** Sangbhin Lee, Jiyeon Hwang, Young-Hwan Jo

## Abstract

The liver plays a central role in energy balance, glucose homeostasis, and lipid metabolism through neural and humoral pathways. Intriguingly, impaired hepatic lipid metabolism has been also associated with an increased risk of anxiety and depression in rodents and humans. However, the mechanisms by which it affects mood behaviors via neural pathways remain poorly understood. This study investigated whether activation of the liver-brain axis can modulate anxiety-like behavior in mice. Advillin (*Avil*)^CreERT2^; channelrhodopsin-tdTomato mice and wireless optogenetics were used to selectively stimulate *Avil*-positive vagal sensory neurons that innervate the liver in freely moving mice. Acute optogenetic stimulation of their nerves in the liver activated neurons in the nodose ganglia and the dorsal motor nucleus of the vagus, and to a lesser extent, those in the nucleus of the solitary tract (NTS). Behavioral assessments revealed that acute optogenetic stimulation of these liver-innervating vagal sensory nerves increased anxiety-like behavior in male and female mice during open field, elevated plus maze, and light/dark box tests. Retrograde viral tracing revealed that neurons in the NTS sent projections to the locus coeruleus (LC), and optogenetic stimulation of liver-innervating vagal sensory nerves resulted in significant activation of norepinephrine-expressing neurons in the LC. Chemogenetic inhibition of LC norepinephrine (NE) neurons completely abolished the anxiogenic effect of stimulating *Slc6a2*□positive vagal sensory neurons, demonstrating that LC NE neuron activity is essential for this behavioral response. Therefore, these findings reveal a novel liver - NTS - LC circuit that plays a role in the regulation of anxiety-like behavior through vagal sensory neurons. Unlike the traditional top-down neuronal circuits associated with the liver, this newly identified liver-brain axis is essential for regulating not only systemic energy homeostasis but also emotional behaviors.

## Introduction

The brain continuously receives and responds to dynamic feedback from afferent (sensory) visceral signals, which influences its functions ^1^. This process is known as interoception and involves bidirectional signal processing between the brain and internal organs to reflect the internal state of an organism ^2^. Vagal sensory neurons in the vagus nerve ganglia innervate most visceral organs, including liver, lung, heart, and gut ^3-6^. Interestingly, the influence of these interoceptive pathways, especially the gut-brain axis, on mental health has been the subject of extensive research, as the gut microbiome plays a significant role in neurodegenerative diseases, cognitive function, and emotion ^7-9^. Additionally, our recent study has revealed that vagal sensory neurons that innervate the liver are also implicated in regulating mood behavior, particularly anxiety-like behavior, in mice fed a high-fat diet (HFD) ^3^. Given that impaired lipid metabolism in the liver significantly affects psychiatric conditions in humans ^10, 11^, liver-innervating vagal sensory neurons that detect changes in hepatic interoceptive signals might contribute to the onset of mental disorders.

The liver is the largest metabolic organ that responds to the energy needs and surfeits. Namely, the liver plays a crucial role in the production and storage of energy in the form of lipids and carbohydrates. Metabolic interactions between the liver and other visceral organs, such as the gut, pancreas, and adipose tissue, are critical for maintaining the energy balance. Particularly, the hepatic portal vein transports blood rich in nutrients and hormones from the gastrointestinal tract and pancreas ^12^. Our recent findings with anterograde and retrograde viral tracing indicate that advillin (*Avil*)-positive vagal sensory neurons that innervate the liver send their nerve terminals predominantly to the portal vein ^3^. This may suggest that these particular sensory neurons are capable of detecting and responding to interoceptive signal molecules, subsequently conveying this sensory information to the dorsal motor complex (DVC), which includes the area postrema (AP), nucleus tractus solitarius (NTS), and dorsal motor nucleus of the vagus (DMV). Importantly, all these DVC nuclei are also innervated by liver-innervating *Avil*-positive vagal sensory neurons ^3^. The caudal NTS is a primary site that integrates visceral sensory information, which then relays to several other brain regions, including the amygdala, paraventricular hypothalamus, dorsal raphe, and locus coeruleus (LC) ^13, 14^.

The LC contains norepinephrine (NE)-expressing neurons, and optogenetic activation of these NE neurons enhances anxiety-like behavior in mice ^15^. Both acute and chronic stimulation of the vagus nerve increase the activity of neurons in the LC in rodents and humans ^16-19^. The improvement in anxiety-like behavior observed in mice lacking liver-innervating vagal sensory neurons in mice fed HFD ^3^ might be attributed to alterations in the liver - NTS - LC pathway via liver-innervating vagal sensory neurons. To gain better insight into how vagal sensory neurons that innervate the liver affect anxiety-like behavior, we expressed channelrhodopsins in *Avil*-positive sensory neurons and optogenetically stimulated their nerve fibers within the liver with a wireless implantable LED device in freely moving animals. Animal behavioral assessments revealed that optogenetic activation of the liver-innervating vagal sensory nerves enhanced anxiety-like behavior compared to the control group. Importantly, this anxiogenic effect was attributed, in part, to the activation of NE-expressing neurons in the LC.

## Materials and methods

### Ethics statement

All mouse care and experimental procedures were approved by the Institutional Animal Care Research Advisory Committee of the Albert Einstein College of Medicine and were performed in accordance with the guidelines described in the NIH guide for the care and use of laboratory animals. Stereotaxic surgery and viral injections were performed under isoflurane anesthesia.

### Animals

All mouse care and experimental procedures were approved by the Institutional Animal Care Research Advisory Committee of Albert Einstein College of Medicine.

Eight-nine-week-old *Avil*^CreERT2^ (stock # 032027), Ai27 (RCL-hChR2(H134R)/tdTomato)-D (stock#), and *Slc6a2*^Cre^ mice (stock# 037882) were purchased from Jackson Laboratory. We crossbred *Avil*^CreERT2^ mice with Ai27D mice to generate *Avil*^CreERT2^;ChR2-tdTomato. Mice were housed in cages at a controlled temperature (22 °C) with a 12:12 h light-dark cycle and were fed a standard chow diet with water provided *ad libitum*. Mice were administered intraperitoneal injections of 75 mg tamoxifen/kg body weight on five consecutive days. The tamoxifen solution was prepared by dissolving it in a mixture of 90% corn oil and 10% ethanol.

### Viral injection

We used a retrograde adeno-associated virus (AAVrg)-Cre (Addgene #55636-AAVrg, titer; 2.2□×□10^13^ pfu/ml), AAVrg-EF1α-double floxed-hChR2(H134R)-mCherry (Addgene #20297-AAVrg, titer; 1.16□×□10^13^ pfu/ml). *Avil*^CreERT2^ and Slc6a2^Cre^ mice received a total volume of 20 ul (5 ul per site/ 4 different sites) of viral injection into the medial and left lobes of the livers. A Hamilton syringe was used to inject over 40 min. The needle (30 G) was left for an additional 10 min to allow diffusion of the virus within the parenchyma.

To selectively inhibit LC NE neurons, a total volume of 200 nl of AAV8-EF1α-DIO-hM4D(Gi)-GFP (Applied Biological Materials, Inc, titer; 1.0 ×□10^12^ pfu/ml) was bilaterally injected into the LC (AP: - 5.4 mm, DV: -3.0 mm, ML: ±0.8 mm). DREADD agonist 21 (MedChemExpress, 1 mg/kg) or saline was intraperitoneally administered 1 h prior to measurement of behavior test.

### Wireless optogenetics

The wireless optogenetic system (Neurolux) consists of a power distribution control box, three autotuners, three modular boxes, peripheral implantable optogenetic devices with μ-LED of 470 nm, and a laptop with NeuroLux software ^20^. In contrast to the traditional wired optogenetic method, this wireless optogenetic approach enables mice to move freely without physical restraints. This device operates without the use of batteries. To implant a peripheral implantable optogenetic device, the animals were anesthetized and the abdominal region was shaved. A 1 cm incision was made on the skin of the abdomen. Blunt forceps were used to form a subcutaneous pocket, and the main body of the device was inserted into the pocket. A 0.5 cm incision was made in the abdominal muscle directly over the liver to position the μ- LED-containing head of the device. This device included two separate LEDs: one with blue emission (470 nm) adjacent to the targeted tissue to serve as the source for optogenetic stimulation, and the other with red emission (650 nm) just under the skin next to the coil to provide an externally visible signal of system activation ^20^. We optogenetically stimulated liver-innervating *Avil*-positive vagal sensory nerve fibers at a frequency of 10 Hz during the animal behavior tests. The stimulation parameters included a pulse duration of 5 s, with each step lasting 10 ms and a 1-second interval between pulses.

### Immunostaining

To assess if optogenetic stimulation increases the neuronal activity, we optogenetically stimulated ChR2-expressing *Avil*-positive nerve fibers in the livers for five minutes and then sacrificed animals 1 hr post stimulation. The mice were anesthetized with isoflurane (3%) and transcardially perfused with 4% paraformaldehyde. Nodose ganglia and brain samples were post-fixed in 4% paraformaldehyde overnight in a cold room, and then in 30% sucrose the following day. Tissues were sectioned using a cryostat at 16-20 μm. The sections were incubated with 0.3% Triton X-100 at room temperature (RT) for 30 min, and then blocked in PBS buffer containing 10% donkey serum and 0.1% Triton X-100 for 1hr at RT and then incubated with rabbit anti-pS6 (Cell signaling, 2215S), goat anti-ChAT (Millipore, AB144P), rabbit anti-RFP (Rockland, 600-407-379), and mouse anti-TH (Millipore, MAB318) antibodies for overnight at cold room, and then sections were washed three times in PBS and incubated with Alexa 594 anti-rabbit IgG for pS6 and RFP (Jackson immunoresearch, 711-585-152), Alexa 488 anti-goat IgG for ChAT (Jackson immunoresearch, 705-545-147), and Alexa 488 anti-mouse IgG for TH (Abcam, ab150105) for 3 hr at RT. The sections were washed, dried, and mounted using VECTASHIELD medium containing DAPI. Images were acquired using a Leica SP8 confocal microscope. Using the cell count plugin in the Image J software (version FIJI), pS6-positive neurons were counted.

### RNAscope *in situ* hybridization (ISH)

For detecting RNA transcripts, we conducted RNAscope ISH analysis. The sections were washed for 10 min in PBS at room temperature (RT), followed by thermal treatment at 60°C for 30 min and post-fixation in 4% PFA at RT for 15 min. The samples then went dehydration through successive 5-min baths in 50%, 70%, and two rounds of 100% ethanol at RT. Target retrieval performed at 95-100°C for 15 min and then the sections were rinsed in ddH_2_O and dehydrated in 100% ethanol for 3 min at RT before air drying. RNAscope protease applied for 30 min at 40°C in a humidified oven. The sections were incubated with *c-fos* (ACD, 316921-C3), *Slc17a6* (ACD, 319171-C2), and *Dbh* (ACD, 407851-C2) for 2 h in a humidified oven at 40°C. Probe amplification involved three 30-min incubation steps with Amp1, 2, and 3 solutions in a humidified oven at 40°C. Signal development was achieved by incubating the sections with their corresponding Vivid dyes. A LEICA SP8 microscope was used for image acquisition.

### Animal behavior tests

Open-field exploration test: Mice were placed in the center of a chamber measuring 40 cm (length) × 40 cm (width) × 40 cm (height) (Maze Engineers) and allowed to explore the chamber for 5 min freely (500 lx throughout the chamber). The center region was designated as 20 x 20 cm^2^. The EthoVision XT video tracking system (Noldus) was used to record the sessions and analyze the behavior, movement, and activity of the animals. The chamber was wiped with 95% ethanol before the subsequent tests to remove any scent clues ^21^. The total distance traveled and time spent in the center and outer zones of the chamber were measured.

Elevated plus maze test: The maze consists of four arms (two open and two closed arms) in the shape of a plus sign (35 cm arm in length, 5 cm arm in width, and 20 cm wall in height) elevated 60 cm from the ground (Maze Engineers). The mice were placed in the center zone and allowed to explore the maze freely for 5 min (500 lx throughout the arms). The session was recorded, and the number of arm entries and amount of time spent in the open and closed arms were analyzed using the EthoVision XT video tracking system.

Light-dark box test: The box consisted of two chambers (one light chamber, 25 cm in length × 40 cm in depth and one dark, 17.5 cm in length × 40 cm in depth, Maze Engineers). Mice were placed in the center of the light chamber and allowed to freely explore the chamber for 5 min (500 lx in the light chamber). The time spent in each compartment and crossing from one compartment to the other were measured using the video tracking system.

### Food intake measurement

To measure food intake, mice were mildly fasted for 6 h from 8:00 am to 2 pm. We optogenetically stimulated liver-innervating *Avil*-positive vagal sensory nerve fibers at a frequency of 10 Hz for 5 while food (Nutra-Gel diet, S5769-TRAY) was presented.

### Assessment of glucose tolerance

For the GTT, the mice were fasted for 15 h (6:00 pm - 9:00 am). Sterile glucose solution was administered intraperitoneally at a concentration of 2 g/kg (glucose/body weigh) at time zero. Immediately after glucose injection, we optogenetically stimulated liver-innervating *Avil*-positive vagal sensory nerve fibers at a frequency of 10 Hz for 5 min. Blood glucose levels were measured at 15, 30, 60, 90, and 120 min after glucose injection. Blood glucose levels were plotted against time after glucose injection, and the area under the curve (AUC) were calculated and compared between groups.

### Measurement of plasma lipocalin-2 levels

Blood samples from mice with and without optogenetic stimulation were collected from the retroorbital plexus using heparinized capillary tubes. Whole blood samples were centrifuged at 3,000 rpm for 10 min, and plasma was separated and stored at -20 °C until use. Plasma lipocalin-2 concentrations were quantified using two-site sandwich ELISA kit (R&D systems, DY1857-05).

### Statistics

All statistical analyses were performed using GraphPad Prism (v.10). Data for each group are presented as mean□±□SEM. Normality of each dataset was assessed using the Shapiro–Wilk test. Group variances were comparable, as confirmed by inspection of residuals and spread of data distributions. For comparisons between two groups, two tailed unpaired Student’s t tests were used. For behavioral assays involving more than two groups, one way ANOVA was performed followed with Tukey’s multiple comparisons test. Sample sizes were based on prior studies using comparable behavioral and optogenetic paradigms; no formal power calculations were conducted. Animals were randomly allocated to experimental and control groups. Mouse body weight was measured to make sure that there was no weight difference between the groups. Behavioral experiments were performed with the experimenter blinded to mouse conditions. Samples in which post-hoc immunohistochemistry indicated unsuccessful viral transduction or malfunction of the optogenetic device were excluded from all analyses. Statistical significance was defined as p□<□0.05.

## Results

### Optogenetic stimulation of liver-innervating *Avil*-positive nerves activates neurons in the DMV and NTS

We first assessed if optogenetic stimulation of liver-innervating *Avil*-positive nerve fibers using our wireless implantable LED device increases the activity of *Avil*-positive neurons in the nodose ganglia of *Avil*^CreERT2^;ChR2-tdTomato mice (Fig. 1A). A 10 Hz light stimulation of the nerve fibers in the liver with a pulse duration of 5 s, with each step lasting 10 ms and a 1-second interval between pulses for five min, effectively activated a small subset of vagal sensory neurons (Fig. 1B). To quantify neuronal activation induced by our stimulation paradigm, we performed immunostaining for phosphorylated S6 (pS6) and RNAscope ISH for *c*□*fos*, two well□ established markers of neuronal activity. Both assays revealed a significant increase in pS6□ and *c*□*fos* □positive cells in the nodose ganglia of stimulated animals compared with unstimulated controls (Fig. 1C). These results indicate that our wireless optogenetic stimulation can reliably activates vagal sensory neurons.

**Figure 1.**
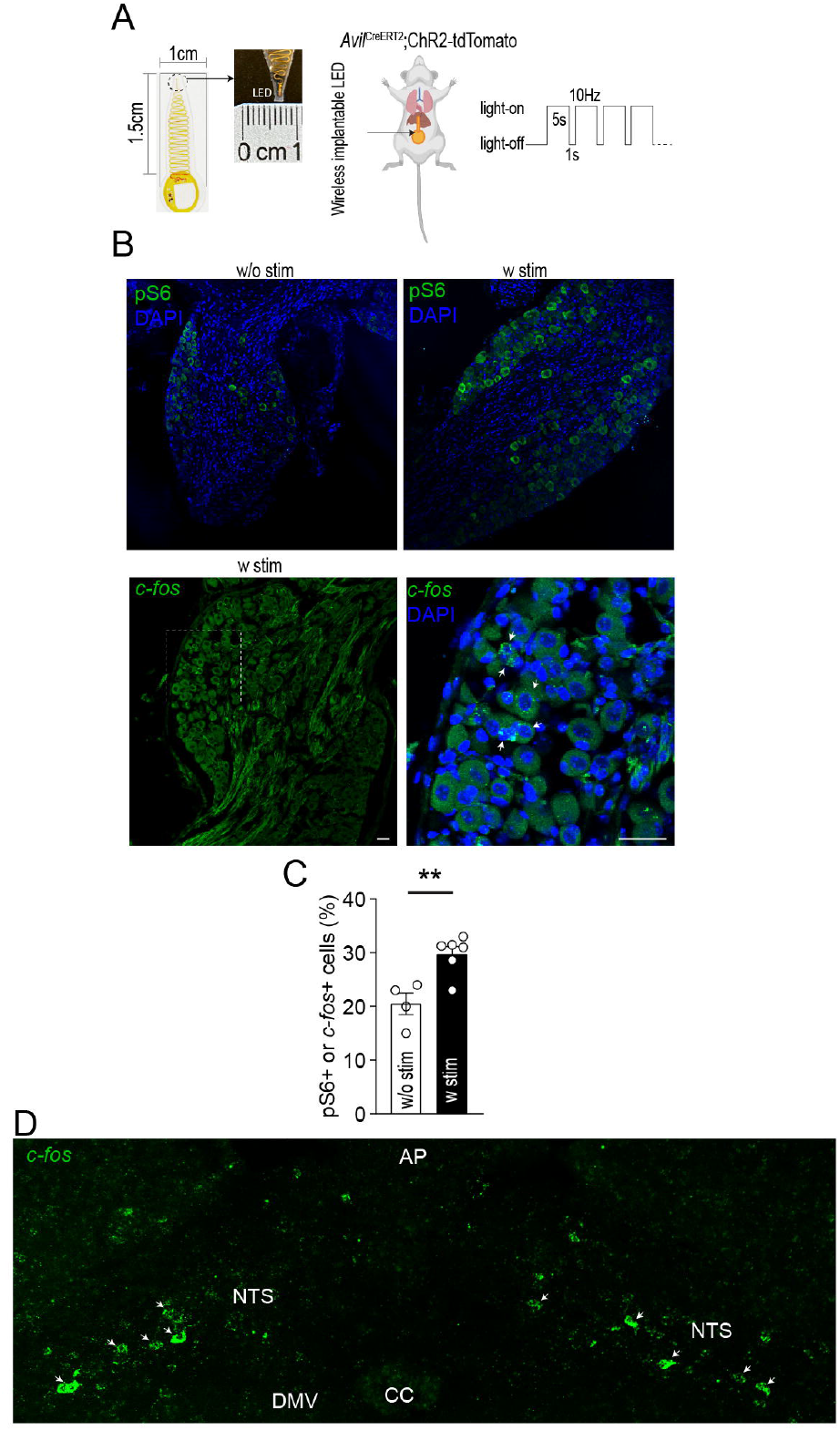
Optogenetic stimulation of liver-innervating *Avil*-positive neurons activates neurons in the NTS. (A) Schematic illustration of the experimental setup. A wireless implantable LED device was positioned directly on the surface of the liver of *Avil*^CreERT2^;ChR2-tdTomato mice. (B) Images of confocal fluorescence microscopy showing pS6-positive neurons and *c-fos*-positive neurons in the nodose ganglia of *Avil*^CreERT2^;ChR2-tdTomato mice, with (right) and without (left) optogenetic stimulation. Scale bar, 40 μm. (C) Quantitation of pS6-positive neurons and *c-fos*-positive neurons in the nodose ganglia (control, n = 4 mice; stimulation, n = 6 mice) revealed a significant increase in activated cells in *Avil*^CreERT2^;ChR2-tdTomato mice with optogenetic stimulation compared to those without it. Unpaired *t*-test, p=0.005. Data are presented as mean ± SEM. (D) Images of confocal fluorescence microscopy showing the expression of *c-fos* in the NTS of *Avil*^CreERT2^;ChR2-tdTomato mice with optogenetic stimulation. White arrowheads indicate the *c-fos*- positive neurons in the NTS. Scale bar, 40 μm.

Because liver□innervating vagal sensory neurons project centrally to the dorsal vagal complex (DVC), we next examine whether activating these fibers is sufficient to drive neuronal activity within this region. Five minutes of optogenetic stimulation of liver□projecting vagal sensory fibers in *Avil*^CreERT2;ChR2□tdTomato^ mice induced *c*□*fos* (Fig. 1D) and pS6 expression (Suppl. Fig. 1A) in a small subset of neurons in the NTS, while eliciting a marked increase in pS6□positive neurons in the DMV (Fig. 1D; Suppl. Fig. 1A). Most pS6-positive cells in the DMV also co-expressed the choline acetyltransferase (ChAT), indicating that they are parasympathetic preganglionic cholinergic neurons (Suppl. Fig. 1A). These findings suggest that both the NTS and the DMV are the downstream targets of liver-innervating vagal sensory neurons.

### Optogenetic stimulation of liver-innervating *Avil*-positive nerve fibers enhances anxiety-like behavior in both male and female mice

We then explored the physiological impact of the acute activation of liver-innervating *Avil*- positive neurons on anxiety-like behavior as we previously showed that liver-innervating vagal sensory neurons influenced anxiety-like behavior in diet-induced obese mice^3^. A series of anxiety-like behavioral assessments was conducted on male mice. First, we performed an open field test, a common assessment for evaluating anxiety-like behavior in rodents^21, 22^. We placed the animals at the center of the open field maze and measured the total distance traveled, along with the time spent and distance traveled in the inner zone. When *Avil*-positive nerves innervating the liver were optogenetically stimulated during the assessment, there was a significant decrease in the total distance traveled compared to that in the control group (Fig. 2A and B), which indicates a reduction in the exploratory locomotor activity during the optogenetic stimulation. Moreover, *Avil*^CreERT2^;ChR2-tdTomato mice with optogenetic stimulation exhibited a decrease in both the distance traveled and the time spent in the inner zone compared to those in the controls (Fig. 2A and B). The percentage of the time spent in the inner zone was also significantly different between the groups (Fig. 2B). However, this anxiogenic like effect was absent in *Avil*^CreERT2;^ChR2□tdTomato mice without light stimulation and in tamoxifen-untreated mice exposed to light stimulation (Suppl. Fig. 2A and B). These findings indicate that acute activation of *Avil*-positive nerves innervating the liver may enhance anxiety-related behavior.

**Figure 2.**
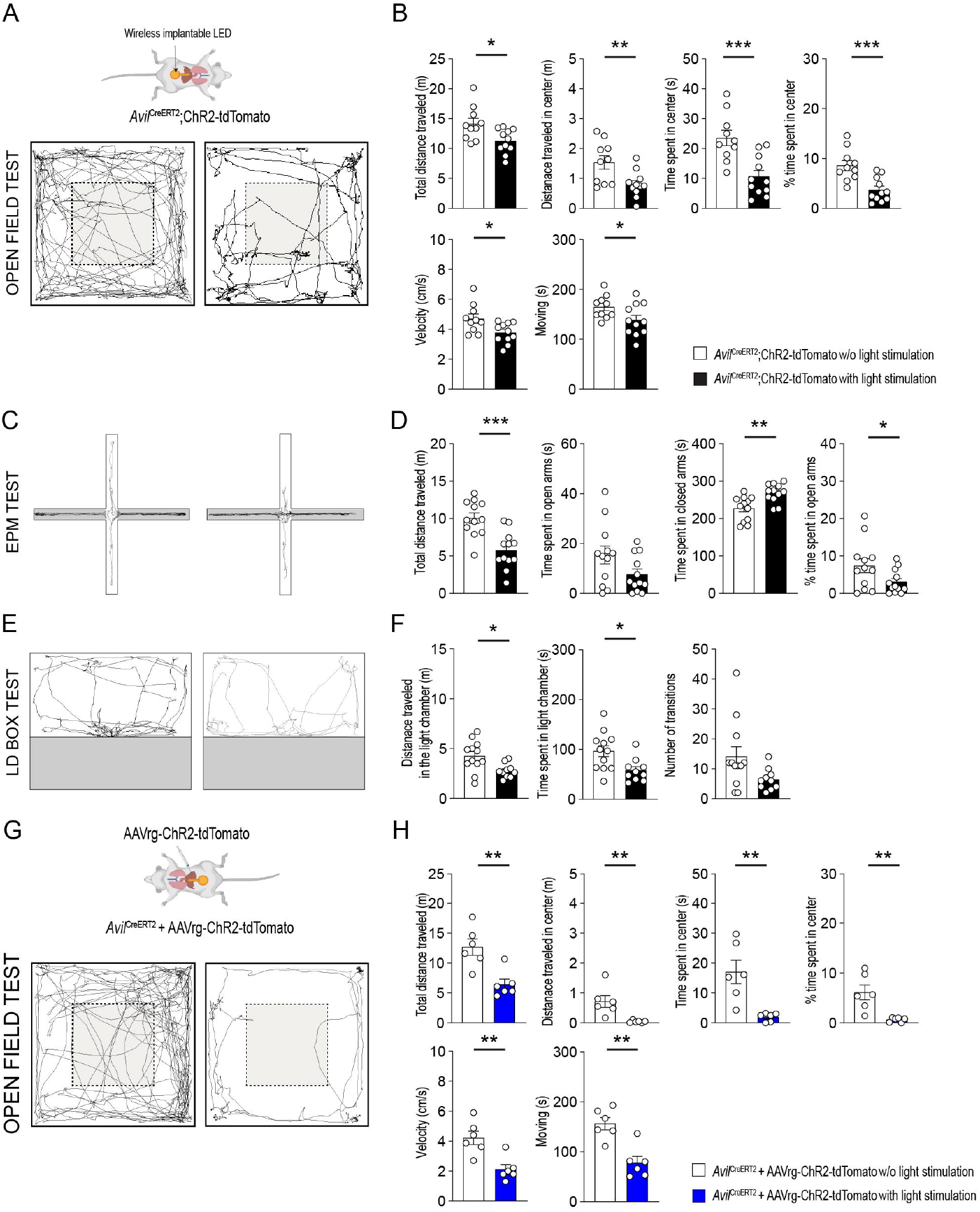
Optogenetic stimulation of liver-innervating *Avil*-positive nerves enhances anxiety-like behavior in male mice. (A and B) Representative examples of the travel paths of mice without (left) and with stimulation (right) during the open field test (A). Stimulation significantly altered total distance traveled (p=0.02), distance traveled in the center (p=0.01), time spent in the center (p<0.001), percentage of time spent in the center (p<0.001), velocity (p=0.02), and moving (p=0.049) were significantly different between the groups (n = 10 mice vs. 11 mice; Two-tailed *t*-test) (C and D) Examples of the travel paths of mice without (left) and with stimulation (right) during the EPM test (C). Stimulation produced significant differences in the total distance traveled (p<0.001), time spent in closed arms (p<0.01), and percentage of time spent in open arms (p=0.045) between the groups (n = 12 mice vs. 12 mice; Two-tailed *t*-test) (E and F) Examples of the travel paths of mice without (left) and with stimulation (right) during the LD box test (E). There were significant differences in the distance traveled in the light chamber (p=0.01) and time spent in the light chamber (p=0.01) between the groups (n = 12 mice vs. 10 mice; Two-tailed *t*-test) (G and H) Representative examples of the travel paths of mice without (left) and with stimulation (right) during the open field test in *Avil*^CreERT2^ mice injected with AAVrg-EF1α-DIO-ChR2-tdTomato into the liver (G). The total distance traveled (p<0.01), distance traveled in the center area (p<0.01), time spent in the center zone (p<0.01), percentage of time spent in the center zone (p<0.01), velocity (p<0.01), and time spent moving (p<0.01) were significantly different between the groups (n = 6 mice vs. 6 mice; Two-tailed *t*-test). Data are presented as mean ± SEM. *p<0.05, **p<0.01, ***p<0.001

We then performed an elevated plus maze (EPM) test to assess the natural exploratory behavior of mice, offering them a choice between the open and enclosed arms. The mice were placed at the center of the EPM. During acute optogenetic stimulation, mice traveled significantly less in the maze than the control group (Fig. 2C and D), further supporting the findings of the open-field test. More importantly, mice with optogenetic stimulation spent less time in the open arms and significantly more time in the closed arms than the controls. The percentage of time spent in the open arms was significantly lower in the mice with stimulation than in those without stimulation. In the light/dark (LD) box test, to evaluate changes in the willingness to explore the illuminated, unprotected area, *Avil*^CreERT2^;ChR2-tdTomato mice with optogenetic stimulation demonstrated a decrease in both the distance traveled and the duration spent in the light chamber compared to the control group (Fig. 2E and F). In addition, stimulation of liver-innervating *Avil*-positive nerves resulted in fewer transitions between compartments. Hence, both the EPM and LD box tests provide further evidence supporting the interpretation that the acute activation of vagal sensory neurons innervating the liver results in an increase in anxiety-like behavior.

We further examined the specificity of *Avil*□expressing vagal afferents to anxiety-like behavior. As in our previous study^3^, we injected a retrograde AAV expressing Cre□dependent ChR2□tdTomato directly into the liver of *Avil*^CreERT2^ mice. Under these experimental conditions, acute optogenetic stimulation of liver-innervating *Avil*-positive sensory nerves consistently enhanced anxiety-like behavior in the open field test, elevated plus maze (EPM) test, and light-dark (LD) box assessment in both male and female mice (Fig 2G-H, and Suppl Fig. 3A-E). These results indicate that liver-projecting *Avil*- positive sensory neurons play a key role in driving anxiety-like responses.

We performed the same behavioral assessments in female mice. The results of the open field test revealed that, similar to male mice, *Avil*^CreERT2^;ChR2-tdTomato mice with acute stimulation exhibited a decrease in the total distance traveled as well as in the time spent and distance traveled within the center zone (Fig. 3A and B). During the assessment of mice on the EPM, acute optogenetic stimulation of their nerves in the liver resulted in a significant reduction in the overall distance traveled and an increase in the time they spent within the closed arms compared to mice that did not receive such stimulation (Fig. 3C and D). In the LD box assessment, female mice exhibited a significant difference in the duration spent in the light chamber and number of transitions between compartments with the presence of optogenetic stimulation (Fig. 3E and F). Moreover, stimulation of liver-innervating *Avil*-positive sensory nerves in *Avil*^CreERT2^ mice injected with AAVrg-ChR2 into the liver also resulted in increased anxiety-like behavior in the open field test, EPM test, and LD box assessment in both male and female mice (Fig 3G-H, and Suppl Fig. 3F-I). Together, these results indicate that acute activation of *Avil*□positive nerves innervating the liver enhances anxiety like behavior in female mice, consistent with the effects observed in males.

**Figure 3.**
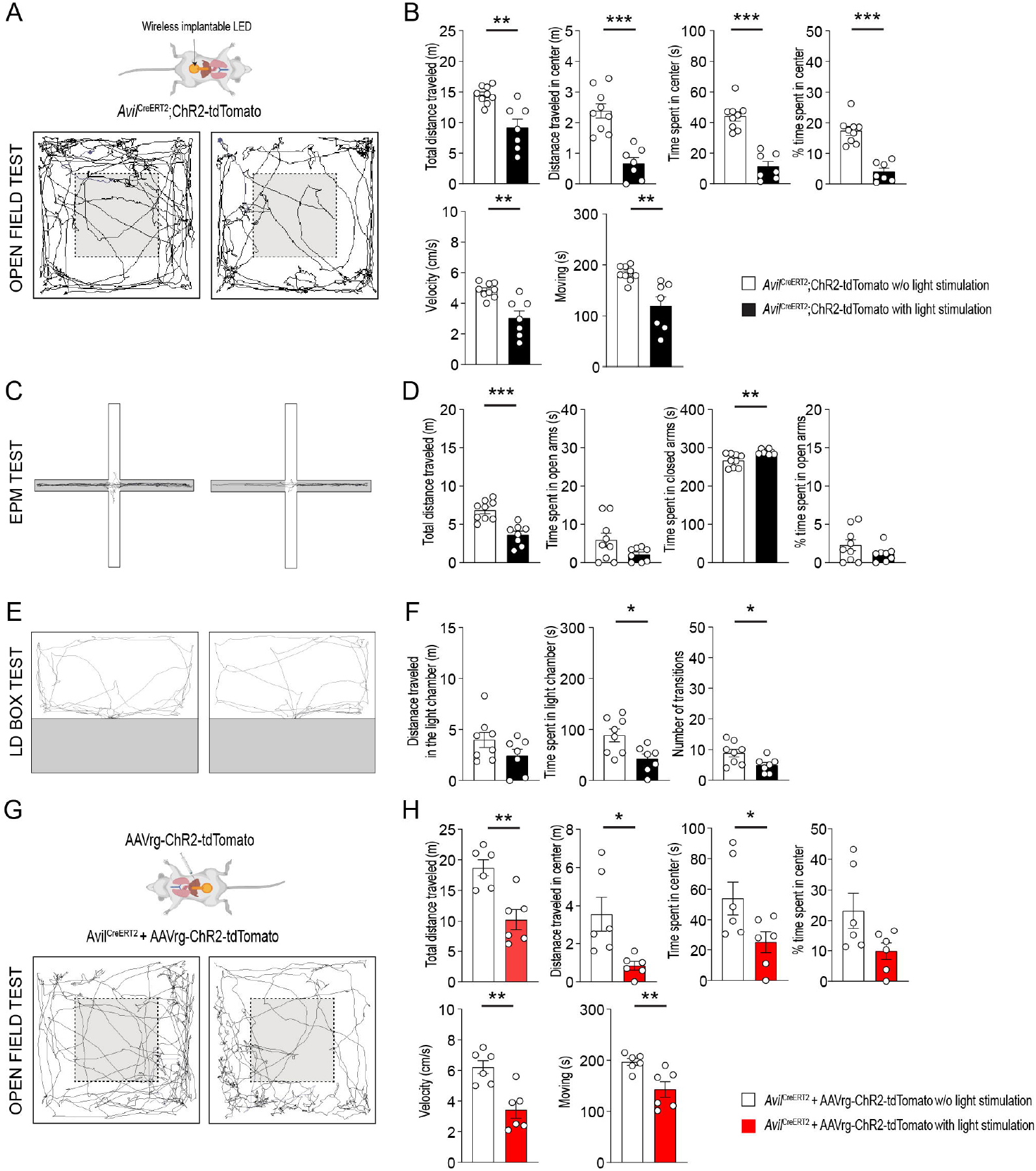
Optogenetic stimulation of liver-innervating *Avil*-positive nerves increases anxiety-like behavior in female mice. (A and B) Representative examples of the travel paths of mice without (left) and with stimulation (right) during the open field test (A). Graphs showing significant differences in the total distance traveled (p<0.01), distance traveled in the center (p<0.001), time spent in the center zone (p<0.001), percentage of time spent in the center zone (p<0.001), velocity (p<0.01), and time spent moving (p<0.01) between the groups (n = 9 mice vs. 7 mice; Two-tailed *t*-test) (C and D) Examples of the travel paths of mice without (left) and with stimulation (right) during the EPM test (C). Significant differences were observed in the total distance traveled (p<0.001) and time spent in the closed arms (p<0.01) between the groups (n = 9 mice vs. 8 mice; Two-tailed *t*-test) (E and F) Examples of the travel paths of mice without (left) and with stimulation (right) during the LD box test (E). Stimulation significantly increased time spent in the light chamber (p=0.01) and number of transitions (p=0.03) (n = 8 vs. 7 mice; Two-tailed *t*-test). (G and H) Representative examples of the travel paths of mice without (left) and with stimulation (right) during the open field test in *Avil*^CreERT2^ mice injected with AAVrg-EF1α-DIO-ChR2-tdTomato into the liver (G). The total distance traveled (p<0.01), distance traveled in the center (p=0.02), time spent in the center zone (p=0.047), velocity (p<0.01), and time spent moving (p<0.01) were significantly different between the groups (n = 6 mice vs. 6 mice; Two-tailed *t*-test). Data are presented as mean ± SEM. *p<0.05, **p<0.01, ***p<0.001

Because this stimulation also activated DMV cholinergic neurons, we next assessed whether it influenced feeding or glucose homeostasis in both male and female mice. Acute optogenetic stimulation did not alter food intake measured immediately after stimulation or 5 minutes later, indicating that short term activation of liver□innervating *Avil*□positive sensory fibers does not acutely affect feeding behavior. (Suppl. Fig. 1B and E). In contrast, during the glucose tolerance test, both male and female mice displayed improved glucose tolerance following stimulation (Suppl. Fig. 1C, D, F, and G). Interestingly, our previous study demonstrated that inhibition of liver innervating cholinergic nerves produced the opposite effect^23^, further supporting the interpretation that altered activity of DMV cholinergic neurons influences systemic glucose handling.

### Optogenetic stimulation of liver-innervating *Avil*-positive nerve fibers activates NE-expressing neurons in the LC

Given that stimulating liver-innervating *Avil*-positive nerve fibers increased anxiety-like behavior, we further explored to understand how this neural pathway triggers such a behavioral response. The LC-NE system plays a crucial role in anxiety-like behavior^24^, and activating the vagus nerve enhances neuronal activity in the LC in both rodents and humans^16-19^. We assessed whether NTS neurons project to LC-NE neurons by injecting AAVrg-tdTomato into the LC of C57BL6/J mice (Fig. 4A). Immunostaining revealed tdTomato-positive cells in the LC, confirming successful viral infection (Fig. 4B). In the same animals, tdTomato-positive cells were also observed predominantly in the NTS (Fig. 4B). Importantly, RNAscope ISH analysis revealed that *c-fos*-positive cells in the NTS co-expressed the excitatory glutamatergic marker *Slc17a6* (Fig. 4C and D), supporting the interpretation that glutamatergic neurons in the NTS are the ones receiving direct synaptic input from liver-innervating vagal sensory neurons. These results may indicate that LC-NE neurons are activated by the stimulation of liver-innervating *Avil*-positive nerves via glutamatergic NTS neurons. Double immunostaining with anti-tyrosine hydroxylase (TH), a marker for catecholaminergic neurons, and anti-pS6 antibodies showed the absence of pS6-positive cells in the LC of unstimulated mice (n = 9 out of 436 TH-positive cells, 2 ± 0.1 %, n= 3 mice; Fig. 4E and H). In contrast, stimulated mice showed robust activation of LC NE neurons, with the majority of TH□positive cells co□expressing pS6 (n = 382 of 505 TH□positive cells; 75□± 1.4%; n□= □5 mice; Fig. 4F and H). Consistent with this, RNAscope ISH analysis demonstrated that optogenetic stimulation induced *c*□*fos* expression in dopamine□β □hydroxylase (*Dbh*)–positive LC neurons (n = 246 of 393 Dbh positive cells; 62□± □4.5%; n□= □3 mice; Fig. 4G, F). Together, these findings support the conclusion that activation of the liver -> NTS -> LC□NE pathway by liver innervating vagal sensory neurons drives LC NE neuron activation, providing a mechanistic basis for the observed increase in anxiety□like behavior.

**Figure 4.**
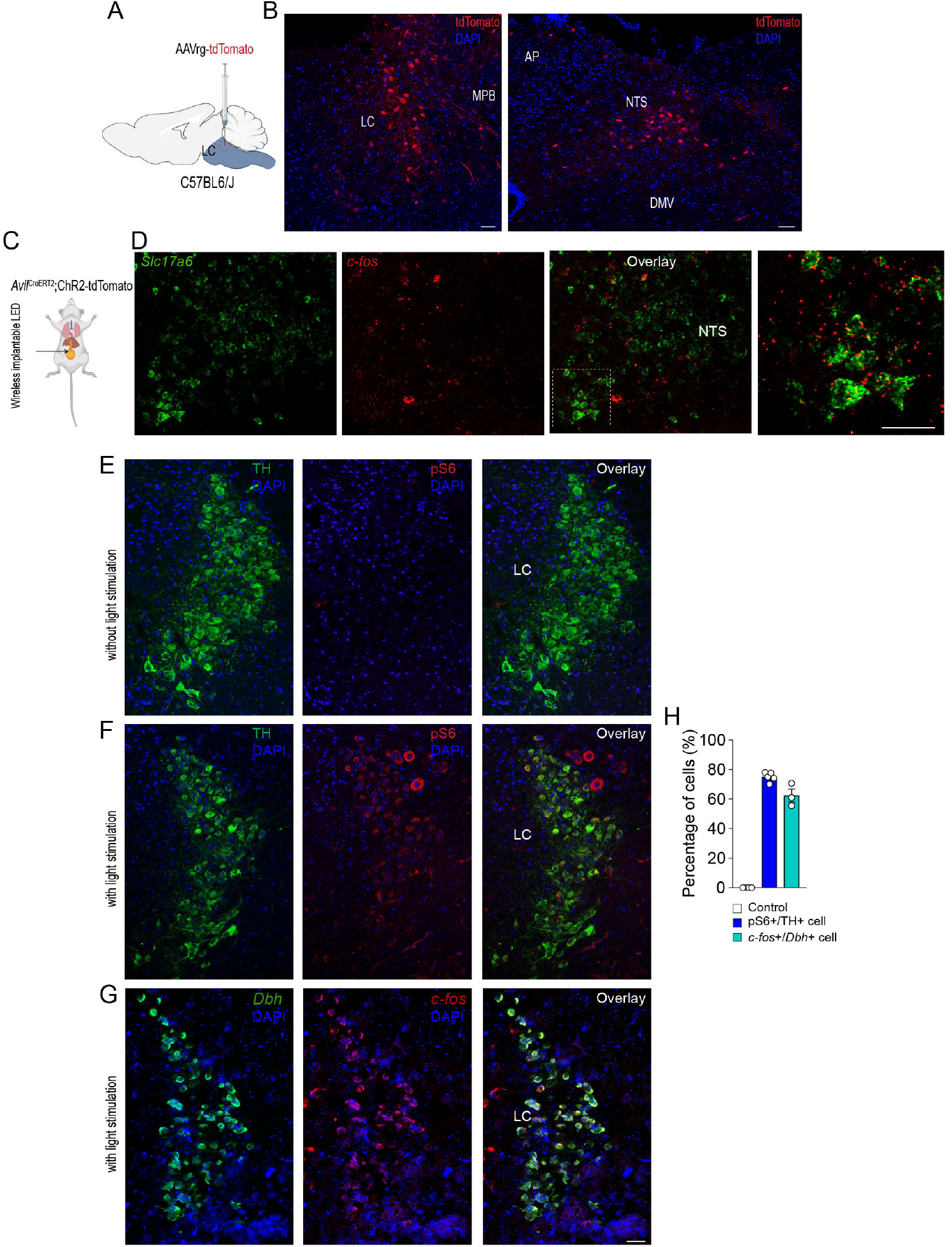
Optogenetic stimulation of liver-innervating *Avil*-positive nerves activates neurons in the LC. (A) Schematic illustration of the experimental design. C57BL6/J mice received stereotaxic infections of AAVrg-tdTomato into the LC. (B) Images of confocal fluorescence microscopy showing tdTomato-positive neurons in the LC of C57BL6/J mice injected with AAVrg-tdTomato, validating successful viral infection. (left) More importantly, tdTomato-positive cells were also detected in the NTS (right), indicating LC-projecting NTS neurons. Scale bar, 40 μm. (C) Schematic illustration of the optogenetic stimulation paradigm. (D) Images from confocal fluorescence microscopy showing *Slc17a6*-positive neurons in the NTS co-expressed *c-fos* in *Avil*^CreERT2^;ChR2-tdTomato mice following optogenetic stimulation. (E). Images from confocal fluorescence microscopy showing that none of the TH-positive neurons in the LC co-expressed pS6 in *Avil*^CreERT2^;ChR2-tdTomato mice without optogenetic stimulation. (F and G) Optogenetic stimulation of liver-innervating *Avil*-positive nerves robustly induced pS6 (F) and *c-fos* (G) expression in the LC. Scale bar, 40 μm (H) Quantification of pS6 and *c-fos*-positive neurons in the LC (Control, n = 4 mice; stimulation, pS6, n = 5 mice, *c-fos*, n = 3 mice) revealed a significant increase in activated neurons in *Avil*^CreERT2^;ChR2-tdTomato mice with optogenetic stimulation compared to those without it.

### Chemogenetic inhibition of LC NE neurons completely blocks the anxiogenic effect of stimulating *Slc6a2*□positive vagal sensory fibers

We further evaluated the functional relevance of the liver - vagal sensory neuron – NTS - LC circuit in regulating anxiety□like behavior. We used solute carrier family 6 member 2 (NE transporter) *Slc6a2*^Cre^ mice, as *Slc6a2* is well-established a neuronal marker for LC NE neurons^25^. Importantly, our previous single□cell RNA□seq analysis of vagal sensory neurons identified 12 transcriptionally distinct clusters, several of which express *Slc6a2* (Fig. 5A). This allowed us to manipulate both a defined subset of liver-innervating vagal sensory neurons and LC NE neurons using the same Cre strain.

**Figure 5.**
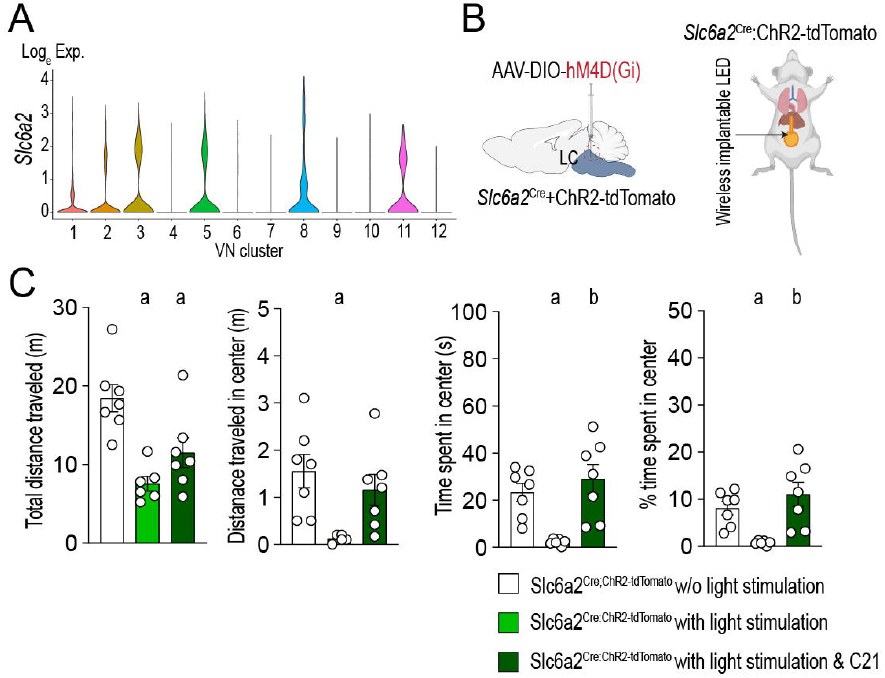
Chemogenetic inhibition of LC NE neurons abolishes the anxiogenic effect induced by activation of liver-innervating *Slc6a2* - positive vagal sensory neurons. (A) Violin plots showing *Slc6a2* gene expression in a subset of vagal sensory neurons (VN). (B) Schematic illustration of the experimental design. *Slc6a2*^Cre^ mice received viral infections of AAV-DIO-hM4D(Gi) into the LC and AAVrg-DIO-ChR2-tdTomato into the liver. (C) Summary of the open filed test assessment across three different conditions. Optogenetic stimulation of liver-innervating Slc6a2-positive nerves significantly reduced distance traveled in the center, time spent in the center, and percentage of time spent in the center test (Control, n□=□7 mice, stimulation, n = 6 mice). In contrast, this anxiogenic effect was completely blocked by inhibition of Slc6a2-positive neurons in the LC (n = 7 mice). One way ANOVA test: a, p<0.05 vs. control (unstimulated); b, p<0.05 vs. stimulated group. Data are presented as mean□±□SEM.

To simultaneously activate peripheral *Slc6a2*□positive vagal sensory fibers and inhibit central LC NE neurons, we injected AAVrg expressing Cre□dependent ChR2 into the liver and AAV encoding Cre□dependent inhibitory hM4D(Gi) into the LC of *Slc6a2*^Cre^ mice (Fig. 5B). Under these experimental conditions, ChR2 was expressed specifically in liver□innervating Slc6a2 positive sensory neurons, while hM4D(Gi) was selectively expressed in LC NE neurons. We optogenetically stimulated liver-innervating *Slc6a2*-positive nerves and measured anxiety-like behavior. This stimulation significantly reduced total distance traveled, distance traveled in center, and time spent in center, indicating an increase in anxiety-like behavior (Fig. 5C). These behavioral changes are consistent with the findings in *Avil*^CreERT2^;ChR2-tdTomato mice.

To directly test whether LC NE neurons are required for this effect, we administered a DREADD agonist one hour before stimulation to selectively silence hM4D(Gi)□expressing LC NE neurons (Fig. 5B). Remarkably, chemogenetic inhibition of LC NE neurons completely abolished the anxiogenic effect of stimulating *Slc6a2*□positive vagal sensory fibers (Fig. 5C), demonstrating that LC NE neuron activity is essential for the behavioral consequences of activating liver□derived vagal sensory input. Together, these findings provide causal evidence that the liver□to□brainstem sensory pathway engages LC NE neurons to modulate anxiety related behavior.

## Discussion

Communication between the visceral organs, particularly the gut and brain has been extensively investigated due to its critical importance in regulating not only energy metabolism such as food intake^6, 26^, energy expenditure^27^, glucose metabolism^3, 28^ but also emotional and mood-related behaviors^29^. As gut-innervating vagal sensory neurons express receptors for nutrients^30^ as well as microbiota-derived metabolites^26^, these interoceptive signal molecules exert influence on these various functions through the gut-brain axis. In addition, the gut-derived interoceptive signal molecules are transported to the liver via the hepatic portal vein that carries blood from other visceral organs, including the gastrointestinal tract, pancreas, and spleen to the liver^12^. Prior studies, including ours, have demonstrated that the vagal afferent nerves sent axonal projections predominantly to the hepatic portal vein^3, 31^. It is plausible that certain functions of the gut-brain axis are also mediated through the liver-brain pathway. In fact, we demonstrated that the loss of function of vagal sensory neurons innervating the liver prevented the onset of diet-induced obesity (DIO) in mice fed a high-fat diet (HFD), partly due to increased energy expenditure^3^. Interestingly, mice without hepatic vagal sensory innervation displayed less anxiety-like behavior during HFD feeding, although their depression-like behavior remained unchanged^3^. These findings imply that disruptions in interoceptive signal detection of liver-innervating vagal sensory neurons may alter mood behaviors, especially anxiety-like behavior.

In our current study, we expanded upon our prior study^3^ to further investigate the critical role of vagal sensory neurons innervating the liver in anxiety-like behavior. Our study specifically examined the acute, neural component of liver□to□brain communication by optogenetically activating liver innervating vagal sensory neurons for a brief period. This approach isolates the fast interoceptive signaling that these neurons normally convey on a moment□to□moment basis during fluctuations in hepatic metabolic state. Our optogenetic stimulation paradigm was therefore intended to mimic this physiological, phasic mode of interoceptive signaling and to determine whether brief activation of this pathway is sufficient to influence anxiety□related behavior. For this, we employed a novel optogenetic stimulation technique that enabled mice to move freely without any physical constraints during the behavioral assessments. This method effectively stimulated the neurons within the nodose ganglia. Despite the relatively small number of pS6□ and *c*□*fos*□positive neurons detected in the NTS after acute stimulation of liver innervating *Avil*□positive fibers, this limited population appears sufficient to drive activation of NE neurons in the LC, consistent with prior studies^16-19^. Supporting this, our retrograde viral tracing showed that LC□projecting neurons were located predominantly within the NTS, and the *c*□*fos*□positive NTS neurons co□expressed the vesicular glutamate transporter, indicating that they are glutamatergic. These findings indicate the existence of a liver -> NTS -> LC NE pathway via liver-innervating vagal sensory neurons. It is well established that directly activating LC NE neurons using optogenetics resulted in enhanced anxiety-like behavior^15^. Hence, our finding of increased anxiety-like behavior following the stimulation of liver-innervating *Avil*-positive neurons would be due in part to the activation of these LC NE neurons. When hepatic vagal sensory innervation was disrupted, the animals exhibited the opposite behavioral phenotype during HFD feeding^3^. Hence, it is plausible that the liver-brain axis is crucial not only for regulating liver functions but also for influencing emotional behavior.

Although our findings highlight the importance of the liver–brain neural circuit in regulating anxiety□like behavior, it is plausible that humoral signals originating from the liver contribute to emotional regulation. In fact, the liver is recognized as an endocrine organ producing and secreting a wide array of hepatokines, cytokines, bile acids, and hormones^32-34^. These circulating factors influence systemic energy metabolism, immune tone, and hepatic function^32, 35, 36^. A recent study demonstrated that stress elevates circulating lipocalin□2 levels in both rodents and humans, and that this cytokine is sufficient to enhance anxiety□like behavior^34^. Intriguingly, restraint stress activated predominantly DMV cholinergic neurons, and optogenetic stimulation of liver□innervating DMV neurons increased lipocalin□2 production in the liver and its release into the bloodstream, thereby promoting anxiety□like behavior^34^.

In our own experiments, acute stimulation of liver-innervating vagal sensory neurons induced pS6 expression in cholinergic neurons in the DMV. A study by Watts et al. using anterograde H129 viruses demonstrated that the DMV cholinergic neurons were the primary central recipients of vagal sensory input from the hepatic portal and superior mesenteric veins without engaging NTS neurons in rats^37^. As acute stimulation of liver-innervating vagal sensory neurons elevated serum lipocalin 2 levels, likely via activation of DMV cholinergic neurons, this raises the possibility that both neural and humoral pathways originating from the liver act in parallel to influence anxiety□like behavior. Specifically, the liver -> vagal sensory neuron -> NTS -> LC circuit may provide a rapid neural route for modulating LC activity, whereas DMV□dependent cytokine release may serve as a slower, reinforcing signal. Hence, future studies will be required to determine whether these pathways interact synergistically to shape the magnitude or duration of anxiety□related responses.

Beyond emotional regulation, our findings also underscore the role of DMV cholinergic neurons in metabolic control. Activation of liver□innervating *Avil* positive neurons did not alter short□term food intake but significantly improved glucose tolerance. Consistent with this, our prior study showed that selective inhibition of liver-innervating parasympathetic cholinergic neurons enhanced hepatic glucose output in mice fed a standard chow diet^23^. Thus, our finding further support the idea that DMV cholinergic neurons are critical regulators of systemic glucose homeostasis. These observations support a model in which a feedforward circuit between the liver and brainstem mediated by vagal sensory neurons and DMV cholinergic neurons coordinates both hepatic energy metabolism and anxiety□like behavior. In addition, loss of liver-innervating vagal sensory neurons reduced the anxiety-like behavior in mice fed HFD in our prior study^38^. Chronic metabolic perturbations such as HFD feeding likely engage slow, adaptive mechanisms - including altered hepatokine secretion, inflammation, and sustained suppression of vagal afferent firing - that reshape the pathway over days to weeks. Thus, acute and chronic manipulations do not represent opposing mechanisms but rather distinct physiological states of the same circuit. Together, these results suggest that the liver - vagal sensory pathway is bidirectionally sensitive to metabolic state, with acute interoceptive signals modulating LC□dependent arousal and anxiety, and chronic metabolic disruption altering this communication through slower, remodeling processes.

In contrast to the traditional view of top□down neural control of liver physiology, our findings reveal a novel bottom□up liver–brain axis that regulates not only systemic energy homeostasis but also emotional behavior. This has important translational implications. Given that steatotic liver disease elevates the risk of anxiety and depression in both rodents and humans^10, 39, 40^, understanding how liver□derived neural and humoral signals shape central neuronal circuits may provide new therapeutic opportunities. Elucidating the mechanisms by which metabolic dysfunction alters this pathway could also help address mood disturbances linked to chronic stress and liver disease.

## References

1. Critchley HD, Harrison NA. Visceral influences on brain and behavior. Neuron 2013; 77(4): 624–638.

2. Chen WG, Schloesser D, Arensdorf AM, Simmons JM, Cui C, Valentino R et al. The Emerging Science of Interoception: Sensing, Integrating, Interpreting, and Regulating Signals within the Self. Trends Neurosci 2021; 44(1): 3–16.

3. Hwang J, Lee S, Okada J, Liu L, Pessin JE, Chua SC et al. Liver-innervating vagal sensory neurons play an indispensable role in the development of hepatic steatosis in mice fed a high-fat diet. Nature communications 2025; 16(1): 991.

4. Chang RB, Strochlic DE, Williams EK, Umans BD, Liberles SD. Vagal Sensory Neuron Subtypes that Differentially Control Breathing. Cell 2015; 161(3): 622–633.

5. Lovelace JW, Ma J, Yadav S, Chhabria K, Shen H, Pang Z et al. Vagal sensory neurons mediate the Bezold-Jarisch reflex and induce syncope. Nature 2023; 623(7986): 387–396.

6. Bai L, Mesgarzadeh S, Ramesh KS, Huey EL, Liu Y, Gray LA et al. Genetic Identification of Vagal Sensory Neurons That Control Feeding. Cell 2019; 179(5): 1129–1143 e1123.

7. Loh JS, Mak WQ, Tan LKS, Ng CX, Chan HH, Yeow SH et al. Microbiota–gut–brain axis and its therapeutic applications in neurodegenerative diseases. Signal Transduction and Targeted Therapy 2024; 9(1).

8. Valles-Colomer M, Falony G, Darzi Y, Tigchelaar EF, Wang J, Tito RY et al. The neuroactive potential of the human gut microbiota in quality of life and depression. Nature Microbiology 2019; 4(4): 623–632.

9. Kolobaric A, Andreescu C, Jašarević E, Hong CH, Roh HW, Cheong JY et al. Gut microbiome predicts cognitive function and depressive symptoms in late life. Molecular Psychiatry 2024; 29(10): 3064–3075.

10. Soto-Angona O, Anmella G, Valdes-Florido MJ, De Uribe-Viloria N, Carvalho AF, Penninx B et al. Non-alcoholic fatty liver disease (NAFLD) as a neglected metabolic companion of psychiatric disorders: common pathways and future approaches. BMC Med 2020; 18(1): 261.

11. Shang Y, Widman L, Hagstrom H. Nonalcoholic Fatty Liver Disease and Risk of Dementia: A Population-Based Cohort Study. Neurology 2022; 99(6): e574–e582.

12. Carneiro C, Brito J, Bilreiro C, Barros M, Bahia C, Santiago I et al. All about portal vein: a pictorial display to anatomy, variants and physiopathology. Insights into Imaging 2019; 10(1).

13. Holt MK. The ins and outs of the caudal nucleus of the solitary tract: An overview of cellular populations and anatomical connections. Journal of neuroendocrinology 2022; 34(6).

14. Kawai Y. Differential Ascending Projections From the Male Rat Caudal Nucleus of the Tractus Solitarius: An Interface Between Local Microcircuits and Global Macrocircuits. Frontiers in Neuroanatomy 2018; 12.

15. McCall JG, Al-Hasani R, Siuda ER, Hong DY, Norris AJ, Ford CP et al. CRH Engagement of the Locus Coeruleus Noradrenergic System Mediates Stress-Induced Anxiety. Neuron 2015; 87(3): 605–620.

16. Groves DA, Bowman EM, Brown VJ. Recordings from the rat locus coeruleus during acute vagal nerve stimulation in the anaesthetised rat. Neurosci Lett 2005; 379(3): 174–179.

17. Frangos E, Ellrich J, Komisaruk BR. Non-invasive Access to the Vagus Nerve Central Projections via Electrical Stimulation of the External Ear: fMRI Evidence in Humans. Brain Stimul 2015; 8(3): 624–636.

18. Dorr AE, Debonnel G. Effect of vagus nerve stimulation on serotonergic and noradrenergic transmission. J Pharmacol Exp Ther 2006; 318(2): 890–898.

19. Hulsey DR, Riley JR, Loerwald KW, Rennaker RL, 2nd, Kilgard MP, Hays SA. Parametric characterization of neural activity in the locus coeruleus in response to vagus nerve stimulation. Exp Neurol 2017; 289: 21–30.

20. Shin G, Gomez AM, Al-Hasani R, Jeong YR, Kim J, Xie Z et al. Flexible Near-Field Wireless Optoelectronics as Subdermal Implants for Broad Applications in Optogenetics. Neuron 2017; 93(3): 509–521 e503.

21. Seibenhener ML, Wooten MC. Use of the Open Field Maze to measure locomotor and anxiety-like behavior in mice. J Vis Exp 2015; (96): e52434.

22. Sestakova N, Puzserova A, Kluknavsky M, Bernatova I. Determination of motor activity and anxiety-related behaviour in rodents: methodological aspects and role of nitric oxide. Interdiscip Toxicol 2013; 6(3): 126–135.

23. Kwon E, Joung HY, Liu SM, Chua SC, Jr., Schwartz GJ, Jo YH. Optogenetic stimulation of the liver-projecting melanocortinergic pathway promotes hepatic glucose production. Nat Commun 2020; 11(1): 6295.

24. McCall JG, Siuda ER, Bhatti DL, Lawson LA, McElligott ZA, Stuber GD et al. Locus coeruleus to basolateral amygdala noradrenergic projections promote anxiety-like behavior. Elife 2017; 6.

25. Caramia M, Romanov RA, Sideromenos S, Hevesi Z, Zhao M, Krasniakova M et al. Neuronal diversity of neuropeptide signaling, including galanin, in the mouse locus coeruleus. Proceedings of the National Academy of Sciences 2023; 120(31).

26. Cook TM, Gavini CK, Jesse J, Aubert G, Gornick E, Bonomo R et al. Vagal neuron expression of the microbiota-derived metabolite receptor, free fatty acid receptor (FFAR3), is necessary for normal feeding behavior. Mol Metab 2021; 54: 101350.

27. Liu C, Angie Lee S, Sun K, Jia L, Lee C et al. PPARγ in Vagal Neurons Regulates High-Fat Diet Induced Thermogenesis. Cell metabolism 2014; 19(4): 722–730.

28. Borgmann D, Ciglieri E, Biglari N, Brandt C, Cremer AL, Backes H et al. Gut-brain communication by distinct sensory neurons differently controls feeding and glucose metabolism. Cell metabolism 2021; 33(7): 1466–1482 e1467.

29. Hung LY, Alves ND, Del Colle A, Talati A, Najjar SA, Bouchard V et al. Intestinal Epithelial Serotonin as a Novel Target for Treating Disorders of Gut-Brain Interaction and Mood. Gastroenterology 2025; 168(4): 754–768.

30. McDougle M, de Araujo A, Singh A, Yang M, Braga I, Paille V et al. Separate gut-brain circuits for fat and sugar reinforcement combine to promote overeating. Cell metabolism 2024; 36(6): 1431.

31. Berthoud HR, Kressel M, Neuhuber WL. An anterograde tracing study of the vagal innervation of rat liver, portal vein and biliary system. Anat Embryol (Berl) 1992; 186(5): 431–442.

32. Jensen-Cody SO, Potthoff MJ. Hepatokines and metabolism: Deciphering communication from the liver. Mol Metab 2021; 44: 101138.

33. Hwang J, Okada J, Liu L, Pessin JE, Schwartz GJ, Jo YH. The development of hepatic steatosis depends on the presence of liver-innervating parasympathetic cholinergic neurons in mice fed a high-fat diet. PLoS Biol 2024; 22(10): e3002865.

34. Yan L, Yang F, Wang Y, Shi L, Wang M, Yang D et al. Stress increases hepatic release of lipocalin 2 which contributes to anxiety-like behavior in mice. Nature Communications 2024; 15(1).

35. Stefan N, Schick F, Birkenfeld AL, Haring HU, White MF. The role of hepatokines in NAFLD. Cell metabolism 2023; 35(2): 236–252.

36. Schwärzler J, Menghini P, Dinarello C, Cominelli F, Tilg H. IL-1 family of cytokines in gastrointestinal and liver disorders. Nature Reviews Gastroenterology & Hepatology 2026; 23(1): 29–43.

37. Garcia-Luna C, Sanchez-Watts G, Arnold M, de Lartigue G, DeWalt N, Langhans W et al. The Medullary Targets of Neurally Conveyed Sensory Information from the Rat Hepatic Portal and Superior Mesenteric Veins. eNeuro 2021; 8(1).

38. Hwang J, Lee S, Okada J, Liu L, Pessin JE, Chua SC, Jr. et al. Liver-innervating vagal sensory neurons are indispensable for the development of hepatic steatosis and anxiety-like behavior in diet-induced obese mice. Nat Commun 2025; 16(1): 991.

39. Labenz C, Huber Y, Michel M, Nagel M, Galle PR, Kostev K et al. Nonalcoholic Fatty Liver Disease Increases the Risk of Anxiety and Depression. Hepatol Commun 2020; 4(9): 1293–1301.

40. Tomiga Y, Tanaka K, Kusuyama J, Takano A, Higaki Y, Anzai K et al. Exercise training ameliorates carbon tetrachloride-induced liver fibrosis and anxiety-like behaviors. Am J Physiol Gastrointest Liver Physiol 2024; 327(6): G850–G860.

