## Supplementary materials for "Optogenetic activation of liver-innervating vagal sensory neurons increases anxiety-like behavior in mice"

Running title: Role of the liver-brain axis in anxiety-like behavior

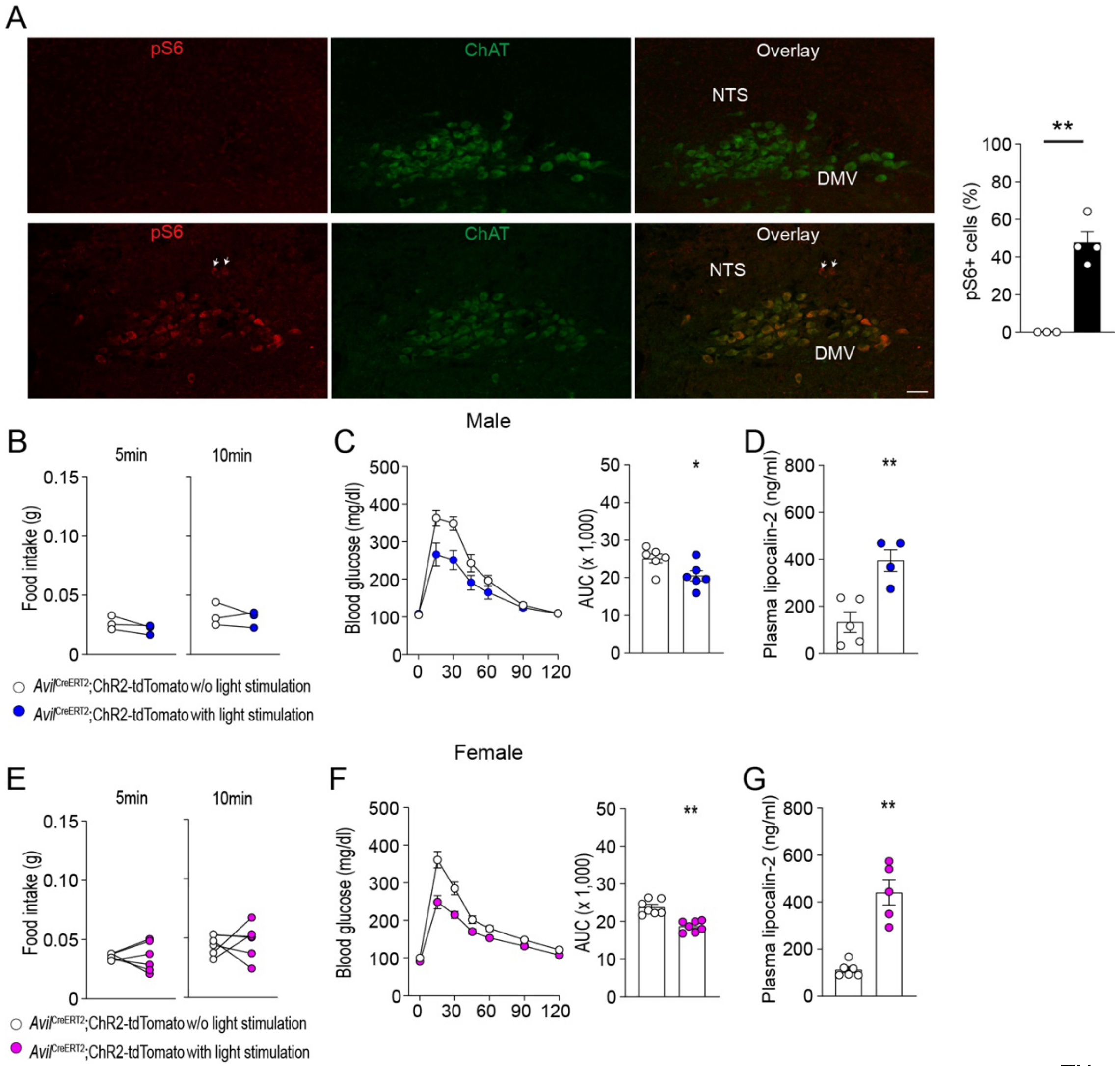


**Supplementary Figure 1**. Optogenetic stimulation of liver-innervating *Avil*-positive neurons activates DMV cholinergic neurons.

(A) Images of confocal fluorescence microscopy showing pS6-positive neurons in the DVC of *Avil*$^{CreERT2}$;ChR2-tdTomato mice without (top) and with (bottom) optogenetic stimulation. White arrowheads indicate the pS6-positive neurons in the NTS. Scale bar, 40 µm. Quantification of pS6-positive neurons in the DMV (n = 3 vs. 4 mice) revealed a significant increase in activated neurons following optogenetic stimulation (Two-tailed t-test, p<0.01).

(B) Graphs showing that food intake in male mice was unchanged by optogenetic stimulation (n = 3 vs. 3 mice).

(C) Summary graphs showing GTT curves for *Avil*$^{CreERT2}$;ChR2-tdTomato male mice with (blue-filled circles) and without (open circles) optogenetic stimulation (n = 6 vs. 6 mice). Area under the curve (AUC) analysis showed a significant difference between groups (Paired t-test, p=0.01).

(D) Graphs showing plasma lipocalin-2 levels in *Avil*$^{CreERT2}$;ChR2-tdTomato male mice with and without optogenetic stimulation (n = 5 vs. 4 mice). Two-tailed t-test, $p<0.01$.

(E) Graphs showing no difference in food intake in females between the groups (n = 6 vs. 6 mice).

(F) Summary graphs showing GTT curves for *Avil*$^{CreERT2}$;ChR2-tdTomato female mice with and without optogenetic stimulation (n = 7 vs. 7 mice). AUC analysis revealed a significant difference between groups (Paired t-test, $p<0.01$).

(G) Graphs showing plasma lipocalin-2 levels in *Avil*$^{CreERT2}$;ChR2-tdTomato female mice with and without optogenetic stimulation (n = 6 vs. 5 mice; Two-tailed t-test, $p<0.001$).

Data are presented as mean ± SEM. *$p<0.05$, **$p<0.01$, ***$p<0.001$

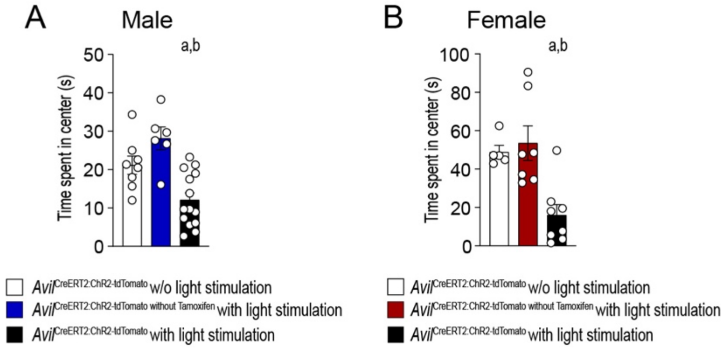


**Supplementary Figure 2**. Optogenetic stimulation of liver-innervating *Avil*-positive nerves enhances anxiety-like behavior in male and female mice.

(A) Graphs showing differences in the time spent in the center in males across three different experimental conditions (*Avil*$^{CreERT2}$;ChR2-tdTomato mice without stimulation, n = 8 mice; tamoxifen-untreated *Avil*$^{CreERT2}$;ChR2-tdTomato mice with stimulation, n = 6 mice; *Avil*$^{CreERT2}$;ChR2-tdTomato mice with stimulation, n= 14 mice).

(B) Graphs showing differences in the time spent in the center in females across three different experimental conditions (*Avil*$^{CreERT2}$;ChR2-tdTomato mice without stimulation, n = 5 mice; tamoxifen-untreated *Avil*$^{CreERT2}$;ChR2-tdTomato mice with stimulation, n = 7 mice; *Avil*$^{CreERT2}$;ChR2-tdTomato mice with stimulation, n= 8 mice).

One-way ANOVA test. a, $p<0.05$ vs. the unstimulated group. b, $p<0.05$ vs. the tamoxifen-untreated stimulation group. Data are presented as mean ± SEM.

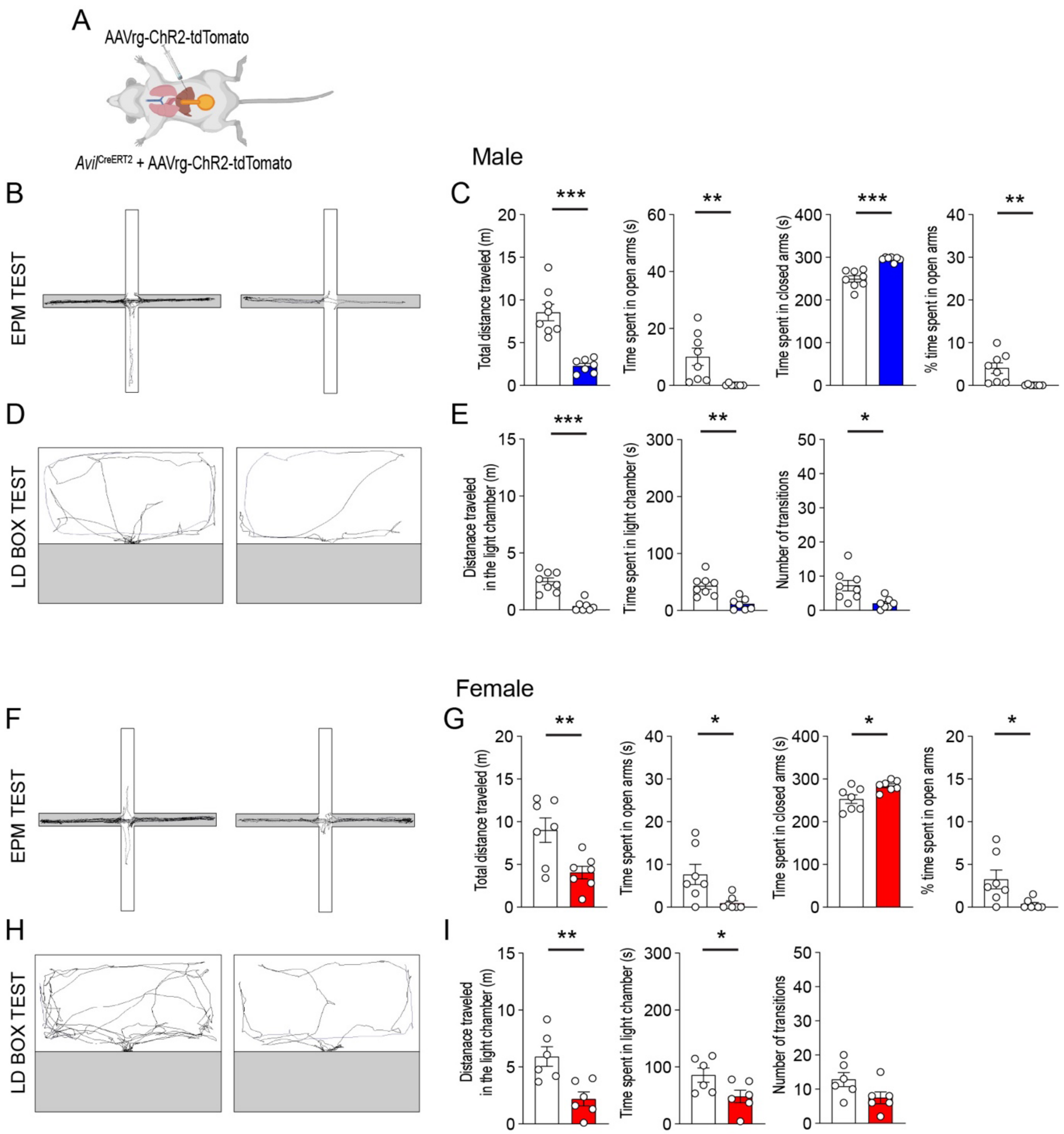


**Supplementary Figure 3**. Optogenetic stimulation of liver-innervating *Avil*-positive nerves increases anxiety-like behavior in male and female mice injected with AAVrg-ChR2 into the liver.

(A) Schematic illustration of the experimental configuration.

(B, C) Representative travel paths of male mice without (left) or with (right) optogenetic stimulation during the EPM test (B). Stimulation significantly altered total distance traveled ($p < 0.001$), time spent in open arms ($p < 0.01$), time spent in closed arms ($p < 0.001$), and percentage of time spent in open arms ($p = 0.01$) ($n = 8$ vs. 7 mice; two-tailed t-test).

(D, E) Representative travel paths of male mice without (left) or with (right) stimulation during the LD box test (D). Stimulation significantly increased distance traveled in the light chamber ($p < 0.001$), time spent in the light chamber ($p < 0.01$), and number of transitions ($p = 0.01$) ($n = 8$ vs. 7 mice; two-tailed t-test).

(F, G) Representative travel paths of female mice without (left) or with (right) stimulation during the EPM test (F). Significant differences were observed in total distance traveled ($p < 0.01$), time spent in open arms ($p = 0.02$), time spent in closed arms ($p = 0.02$), and percentage of time spent in open arms ($p = 0.02$) ($n = 7$ vs. 7 mice; two-tailed t-test).

(H, I) Representative travel paths of female mice without (left) or with (right) stimulation during the LD box test (H). Stimulation significantly increased distance traveled in the light chamber ($p < 0.01$) and time spent in the light chamber ($p = 0.047$) ($n = 6$ vs. 6 mice; two-tailed t-test). *$p<0.05$, **$p<0.01$, ***$p<0.001$

### Abbreviations

AAV, adeno-associated virus

AP, area postrema

AUC, area under the curve

Avil, Advillin

ChAT, choline acetyltransferase

ChR2, channelrhodopsin-2

Dbh, dopamine-beta-hydroxylase

DIO, diet-induced obesity

DMV, dorsal motor nucleus of the vagus

DVC, dorsal motor complex

ELISA, enzyme-linked immunosorbent assay

EPM, elevated plus maze

GTT, glucose tolerance test

HFD, high-fat diet

LC, locus coeruleus

LD, light/dark

NE, norepinephrine

NTS, nucleus of the solitary tract

pS6, phospho-S6 ribosomal protein

RT, room temperature

TH, tyrosine hydroxylase
